# High-speed Auto-Polarization Synchronization Modulation Three-dimensional Structured Illumination Microscopy

**DOI:** 10.1101/2023.12.04.569876

**Authors:** Yaning Li, Ruijie Cao, Wei Ren, Yunzhe Fu, Yiwei Hou, Suyi Zhong, Karl Zhanghao, Meiqi Li, Peng Xi

## Abstract

In recent years, notable progress has been achieved in both the hardware and algorithms of structured illumination microscopy (SIM). Nevertheless, the advancement of 3DSIM has been impeded by challenges arising from the speed and intricacy of polarization modulation. In this study, we introduce a high-speed modulation 3DSIM system, leveraging the polarization maintaining and modulation capabilities of a digital micro-mirror device (DMD) in conjunction with an electro-optic modulator. The DMD-3DSIM system yields a 2-fold enhancement in both lateral (133 nm) and axial (300 nm) resolution compared to wide-field imaging, and can acquire a data set comprising 29 sections of 1024×1024 pixels, with 15 ms exposure time and 6.75 s per volume. The versatility of the DMD-3DSIM approach was exemplified through the imaging of various specimens, including fluorescent beads, nuclear pores, microtubules, actin filaments, and mitochondria within cells, as well as plant and animal tissues. Notably, polarized 3DSIM elucidated the orientation of actin filaments. Furthermore, the implementation of diverse deconvolution algorithms further enhances three-dimensional resolution. The DMD-based 3DSIM system presents a rapid and reliable methodology for investigating biomedical phenomena, boasting capabilities encompassing 3D superresolution, fast temporal resolution, and polarization imaging.

## 1 Introduction

The 2014 Nobel Prize in Chemistry is awarded for the groundbreaking advancement of superresolution microscopy, which surpasses the fundamental diffraction limit of spatial resolution^1-4^. Among these remarkable technologies, structured illumination microscopy (SIM) stands out as a unique and highly compatible approach that can provide ultra-high temporal and spatial resolutions. SIM can be mainly categorized into two types: two-dimensional structured illumination microscopy (2DSIM)^5-8^ and three-dimensional structured illumination microscopy (3DSIM)^9-11^. 2DSIM, also known as traditional SIM, extends the resolution beyond the diffraction limit by a factor of 2 in the *xoy* plane by utilizing 2D patterned illumination and subsequent reconstruction algorithms^7, 12-16^ 2DSIM has emerged as a powerful imaging technique that facilitates rapid and time-lapse visualization of diverse biological processes. However, due to the illumination pattern of 2DSIMis not modulated axially, its axial resolution is the same as that of wide-field microscopy, leading to significant reconstruction errors when imaging thick specimens with defocus background. In stark contrast, 3DSIM with axial-modulated illumination can effectively double the axial resolution. This enables super-resolution imaging in all three dimensions (*xyz*), enhancing the discernment of subcellular structures with unparalleled clarity and accuracy, particularly in deeper regions of the specimen. And unlike traditional 2DSIM images^17, 18^, which are flat and lack depth information, three-dimensional imaging aims to recreate the spatial characteristics of the subject, providing a more realistic and immersive representation.

Various implementations have been developed to realize the structured illumination and polarization modulation of the 3DSIM system, whose crucial device includes phase grating^9, 19^ and ferroelectric liquid crystal spatial light modulator (FLC-SLM). Phase grating^5, 9, 20^ with a fixed linear polarizer is rotating and laterally translated to control the direction, phase, and polarization of the structured light, but the imaging speed is limited by the mechanical rotation (∼1 s) and transversal movement (∼10 ms) of the phase grating, and the incompatibility of phase grating period and interference frequency at different excitation wavelengths poses a challenge for multicolor imaging. FLC-SLM applies electric fields to the crystal layer and realizes the modulation of the optical field, which plays a crucial role by acting as a digital grating, replacing the traditional transmission phase grating. However, FLC-SLM cannot continuously display patterns, which requires constant switching of images and precise timing synchronization. And then, the response time of ferroelectric liquid crystal phase retarders is normally milliseconds, which will extremely drag the imaging speed and cause motion artifacts. What’s more, Pizza wave plates are unable to adjust the zeroth-order light, necessitating a separate light path for its introduction and separate adjustment of its polarization angle when using SLMs, posing the difficulty to construct SLM-based 3DSIM system.

The digital micro-mirror device (DMD) is a kind of spatial light modulator that utilizes the electro-mechanical rotation of micromirrors to modulate the light field reflecting off DMD. Its fast response speed, low cost, and wide availability make it a popular choice for projectors and scientific research instruments. As each micro-mirror is controlled in the binary form corresponding to on and off states, DMD can also become a digital reflection grating when loading with a specific pattern and thus can provide a rapid switch of structured illumination patterns for 3DSIM. Compared with FLC-SLM-based 3DSIM, DMD-based 3DSIM possesses the following advantages in the interference SIM system. First, after loading pattern images, DMD maintains in working state and does not require a refresh cycle, as FLC-SLM does, which greatly simplifies the timing control of the SIM system; Then, DMD has a higher switching speed, with the latest products capable of switching speeds up to 23 kHz/1 bit (DLP9500, TI), making it more suitable for fast imaging of live cells. Additionally, due to the nature of the coating on the DMD surface, DMD can preserve the polarization state of reflected light the same as the incident light. With an electro-optic modulator (EOM) capable of switching speeds up to the order of nanoseconds, DMD enables ultra-fast imaging with minimal motion artifacts. All of these aspects show the greater potential and inspire us to combine DMD and EOM to build a 3DSIM system. (Supplementary S1 shows the comparison of three representative SIM methods with various polarization control schemes, and Supplementary S2 shows the polarization extinction ratio of DMD-based SIM with three polarization control schemes). Overall, SIM with its remarkable capabilities and flexibility, combined with the distinct advantages offered by DMDs, holds tremendous potential for advancing super-resolution microscopy techniques.

In this work, we present a novel and high-speed auto-polarization synchronization modulation three-dimensional structured illumination microscopy system, leveraging the auto-polarization-maintenance capability of DMD, in conjunction with the polarization modulation capability of electro-optic modulators (EOM) for the first time. By harnessing these advanced technologies, our developed system demonstrates exceptional super-resolution imaging capabilities in all three dimensions (*xyz)* and imaging speed, while enabling the precise determination of fluorescent dipole orientations. Through extensive experimentation, we successfully imaged a diverse array of subcellular structures, including the nuclear pore complex, microtubules, actin filaments, and mitochondria in animal cells. Moreover, we extended the application of our 3DSIM system to investigate highly scattering plant cell ultrastructures, examining cell walls in oleander leaves, hollow structures in black algal leaves, and features within the root tips of corn tassels. Notably, in a mouse kidney slice, the actin filaments exhibited a pronounced polarization effect, which was effectively captured and visualized using our 3DSIM approach. Overall, our innovative DMD-based 3DSIMtechnique offers a rapid, precise, and versatile platform for super-resolution imaging, facilitating a range of significant biological discoveries and providing a reliable foundation for the advancement of next-generation 3DSIM.

## 2 Methods

### 2.1 Optical Setup

An optical schematic for the 3DSIM setup is illustrated in Fig. 1(a). An excitation laser beam of 561nm was polarized via a Glan-Taylor polarizing prism and modulated by a polarization rotator composed of an electro-optic modulator and a quarter wave plate. With linearly polarized light incident on the polarization rotator, output polarization can be rotated in azimuth at a speed up to 250 kHz by adjusting the liquid crystal retardation of EOM. The emergent light was collimated and expanded to a diameter of 13 mm via a beam expander and then illuminated on the pattern generator DMD, a 1920×1080-pixel spatial light modulator, at an incidence angle of 12° directed by mirrors. Each pixel of DMD can tilt along the diagonal of the square micromirror and the tilt direction of each micromirror is dictated by the binary contents of the CMOS memory cell associated with each micromirror. A binary value of 1 results in a micromirror landing in the ON State direction and a binary value of 0 results in a micromirror landing in the OFF State direction. Therefore, when a specific periodic binary fringe is loaded on the DMD, the DMD works as a pattern generator, and the diffracted beams converging through lens1 can form 0^th^ and ±1 ^st^ order beams perpendicular to the periodic binary fringe. The custom-built spatial mask consisting of one central 0.6 mm diameter pinhole with an attenuation filter and six symmetrical 0.6 mm diameter pinholes is located in a pupil plane of L1 and can screen out the wanted seven diffraction beams and adjust the intensity ratio of 0^th^ and ±1 ^st^ beams. The intensity of order 0 was 75% of that of ±1^st^ orders to strengthen the highest-lateral-frequency information components. The 0^th^ and ±1 ^st^ order diffraction beams were refocused to the center and two points near the opposite edges of the back focal plane of the objective lens through a 4f system consisting of lens2 and lens3 via a polarization-preserving dichroic mirror coating with minimal transmission retardance at the excitation wavelength to maintain the desired polarization state of the illumination beam^21^. The objective recollimated the three beams and made them approach the sample and intersect with each other to produce a 3D pattern of excitation intensity with both axial and lateral structures in the focal plane of the objective. The fluorescence emitted from the sample was gathered by the same objective and passed through the same polarization-preserving dichroic mirror, a tube lens, and an emission filter to a water-cooled sCMOS camera with 82% peak quantum efficiency to detect the emission fluorescence. The follow-up instrument is discussed in Supplementary S3.

**Fig. 1.**
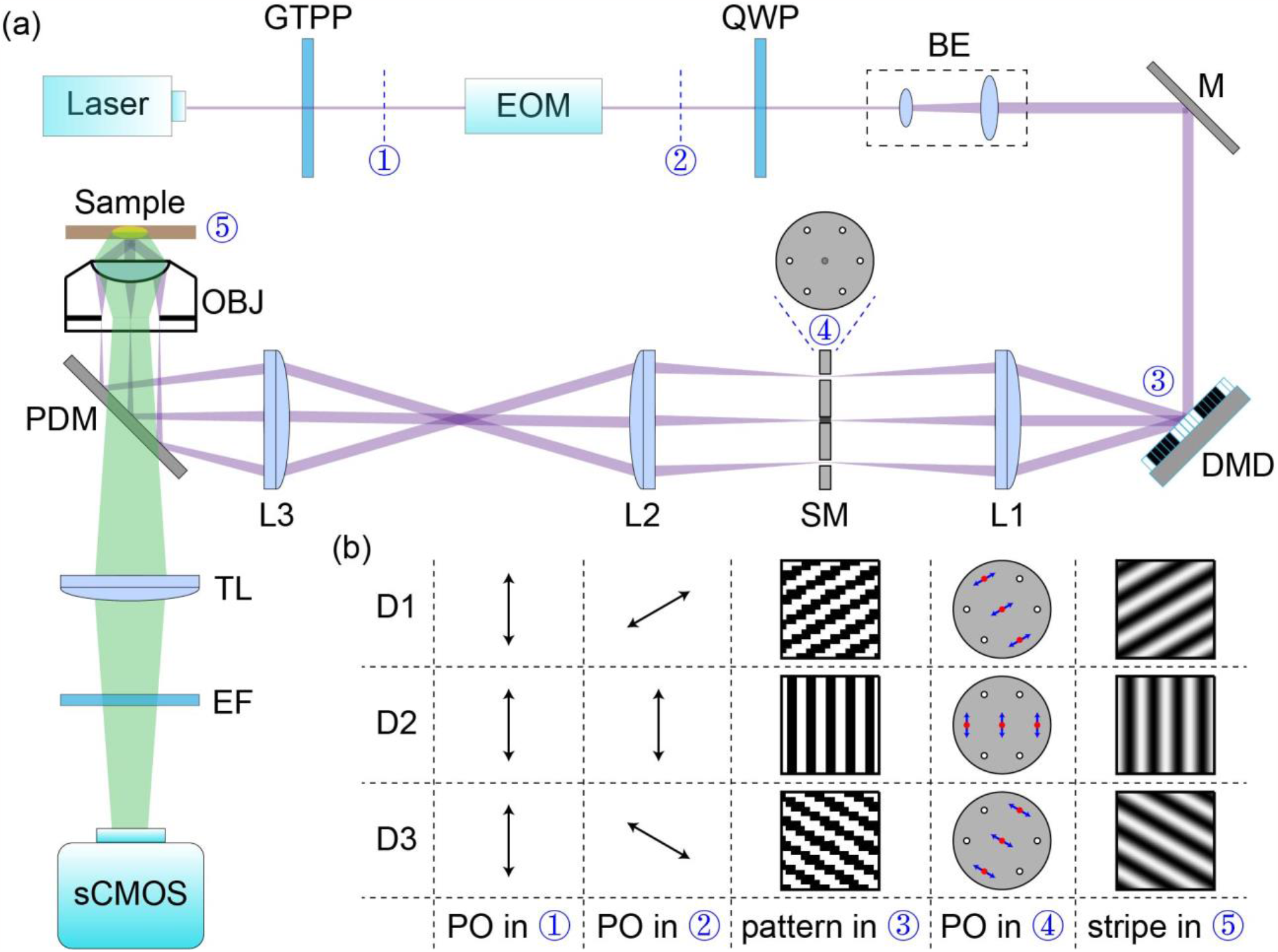
Diagrams of the hardware of 3DSIM. (a) Optical setup. (b) The illumination polarization before entering the EOM remains the same (column 1) and the beam that passes through the polarization rotator exhibits different polarization states (column 2) due to the property of linear polarization rotation of the polarization rotator. Three patterns with different orientations were preloaded in DMD (column 3). The positions of the 0 and ±1 order beams converged to the SM (column 4). Three Interference stripes with different orientations in the sample (column 5). GTPP, Glan-Taylor polarizing prism; EOM, electro-optic modulator; QWP, quarter wave plate; BE, beam expander; M, mirrors; L, lens; SM, spatial mask; OBJ, objective lens; PDM, polarization-preserving dichroic mirror; TL, tube lens; EF, emission filter.

### 2.2 Data Acquisition and Time Sequence of the Synchronization of the System

The acquisition of three-dimensional data involved the use of five pattern phases spaced evenly at intervals of 2π/5, three pattern orientations separated by 60 degrees, and an axial step size of 125 nm. The phase and orientation of the illumination stripe were controlled by rotating and laterally translating the pattern in DMD. When patterns in Fig. 1(b) were preloaded onto the DMD controller circuit board, DMD served as a grating, enabling precise and reproducible control of the position of the stripes formed by interference in the sample along the direction of its pattern wave vectors under the accurate determination of pattern wave vectors. We employed a piezoelectric translator (NanoMax 300, Thorlabs) controlled by closed-loop feedback from a capacitive position sensor to allow axial movement of the sample holder to the objective lens. And, the acquired data spanned an axial range that typically extended slightly above and below the region of interest. In certain cases where special requirements for high-speed imaging exist, sCMOS cameras can be operated in the rolling shutter mode (synchronous readout trigger mode), where exposure is triggered at the beginning of each frame and progresses row by row until the entire frame is captured. The synchronization of optical system components is crucial for proper functioning and performance. In our system, the differential amplifier (Model 302RM, Conoptics) of EOM for polarization rotating, DMD pattern switching and displaying, the trigger for the sCMOS camera to exposure in synchronous read-out mode, and trigger for the piezoelectric translator to move along axial direction were all precisely synchronized by a LabVIEW-based program running on a data acquisition equipment (PCIe-6738, National Instruments). The time sequence of the synchronization of DMD-based 3DSIM is in Supplementary S4.

### 2.3 Illumination and Reconstruction Principle of 3DSIM

Because of the characteristics of the optical system, the optical transfer function (OTF) has a leaky-cone shape^9^. This causes a defocus background and limitation in the z-axis. Through shifted spectrum to fill the OTF, 3DSIMachieves the double enhancement in the *xyz* axis as shown in Fig. 2(a, b). When the three beams interface in the specimen plane ***S(x, y, z)***, the emission light distribution ***C***_*θ,φ*_***(x, y, z)*** with illumination angle *θ* and phase *φ* can be expressed as:

**Fig. 2.**
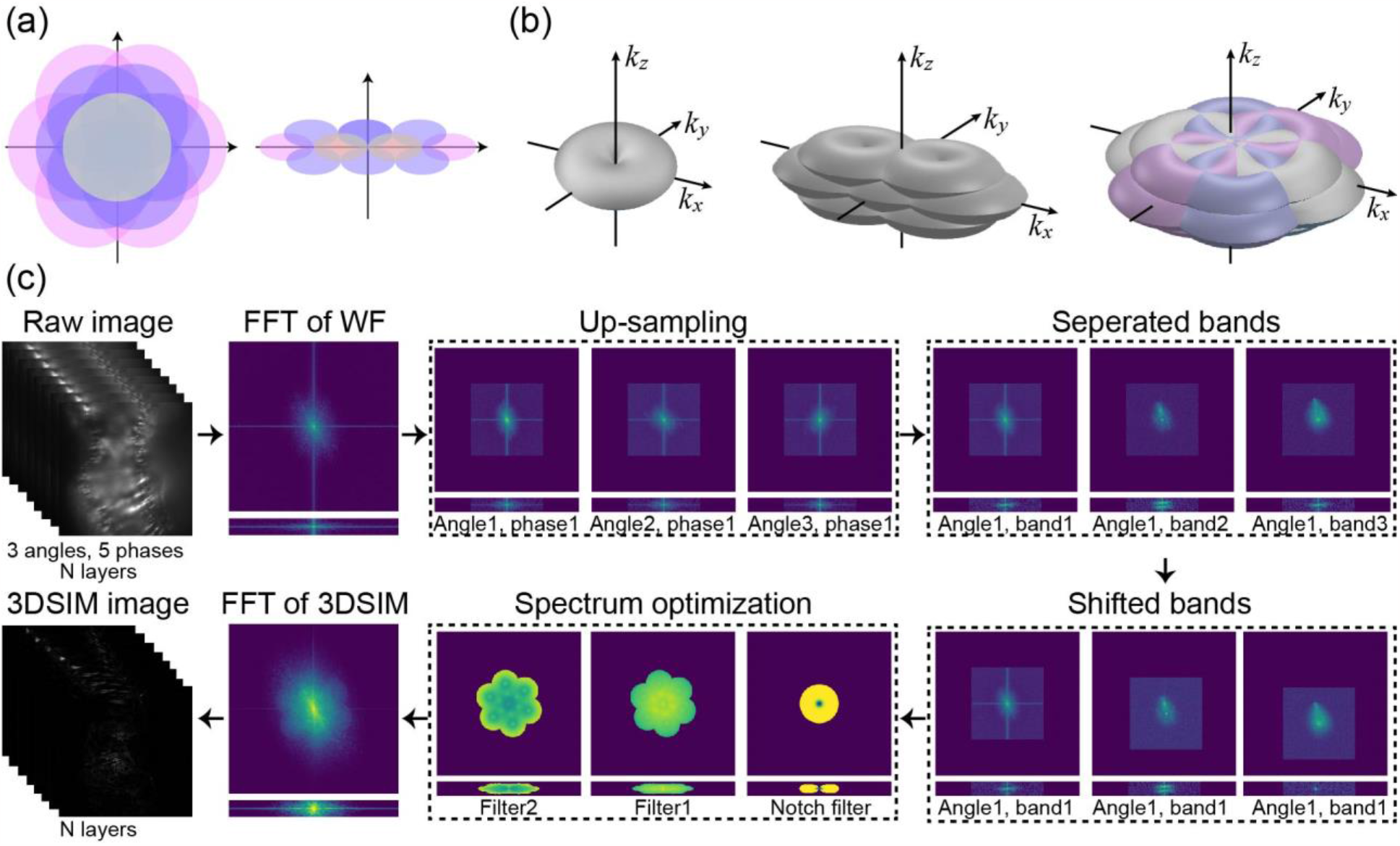
Principle and algorithm flow of 3DSIM. (a) Separated spectrum bands fill the leaky-cone OTF and double the three-dimensional spectrum with the corresponding 3D insight in (b). (c) The algorithm flow of 3DSIM, including the middle spectrum results.

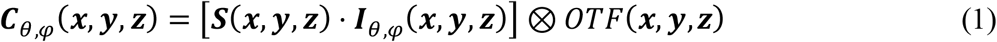

Where ***I***_*θ,φ*_ ***(x, y, z)*** is the excitation illumination light, *OTF****(x, y, z)*** is the leaky-cone OTF of the 3DSIM system.

Different from 2D-SIM, which has 0^th^ and 1^st^ harmonics, the illumination light of 3DSIMcan be composited into the 0^th^, 1^st^, and 2^nd^ harmonics, and can be expressed as^22^:

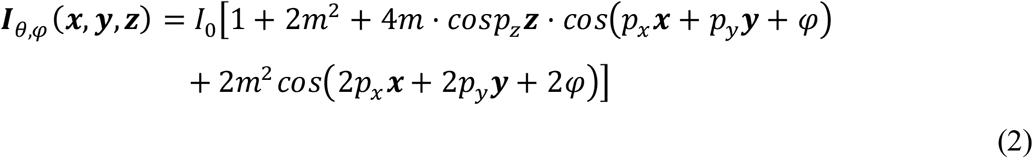

where ***(x, y, z)*** are the vector bases of *xyz* in three-dimensional space. We can see that only the 1^st^ harmonic (4*m* · *cos*2π*p*_*z*_***z***) is relevant to the *z*-axis and the other harmonics (1 + 2*m*^2^ and 2*m*^2^ ) are only relevant to the *xy*-axis. So, let *m*_**0**_ = 1 + 2*m*^2^, *m*_1*z*_***(z)*** = 4*m* · *cos*2π*p*_*z*_***z*** (let *m*_1_ = 4*m*) and *m*_2_ = 2*m*^2^ to simplify the expression. The Fourier transform of illumination light is:

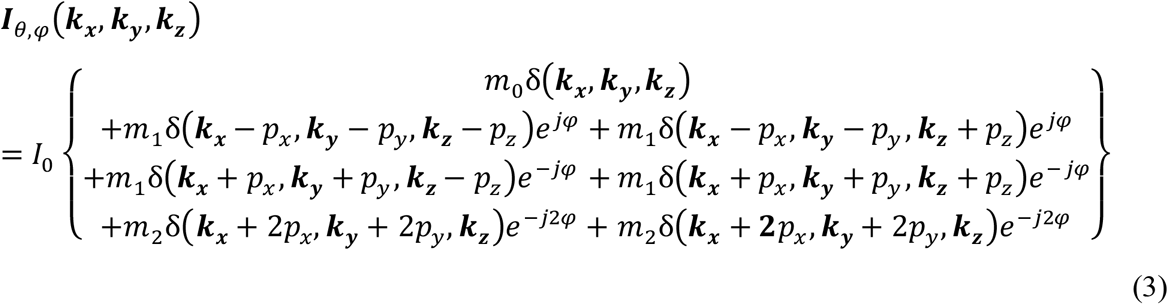

So, the 3D Fourier transform of emission light distribution ***C***_*θ,φ*_***(x, y, z)*** can be expressed as:

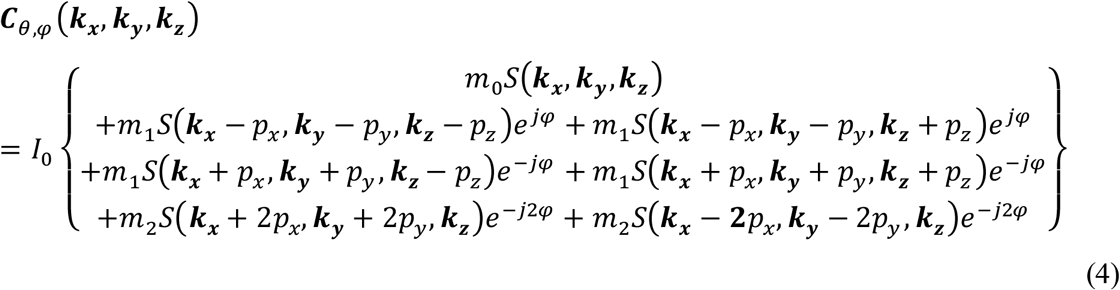

Normally we set the three illumination angles of *θ*_1_, *θ*_2_ = *θ*_1_ + 2π/3, *θ*_3_ = *θ*_1_ + 4π/3 to improve the isotropy of reconstruction. In each illumination angle, we can resolve the 0^th^, ±1^st^, and ±2^nd^ bands (i.e. *S****(k***_***x***_, ***k***_***y***_, ***k***_***z***_), *S****(k***_***x***_ ± *p*_*x*_, ***k***_***y***_ ± *p*_*y*_, ***k***_***z***_ − *p*_*z*_ ) + *S****(k***_***x***_ ± *p*_*x*_, ***k***_***y***_ ± *p*_*y*_, ***k***_***z***_ + *p*_*z*_), and *S****(k***_***x***_ ± 2*p*_*x*_, ***k***_***y***_ ± 2*p*_*y*_, ***k***_***z***_)) through the five images of phase (typically *φ*_1_ = 0, *φ*_2_ = 2π/5, *φ*_3_ =4π/5, *φ*_4_ = 6π/5, *φ*_5_ = 8π/5) using the matrix separation method.

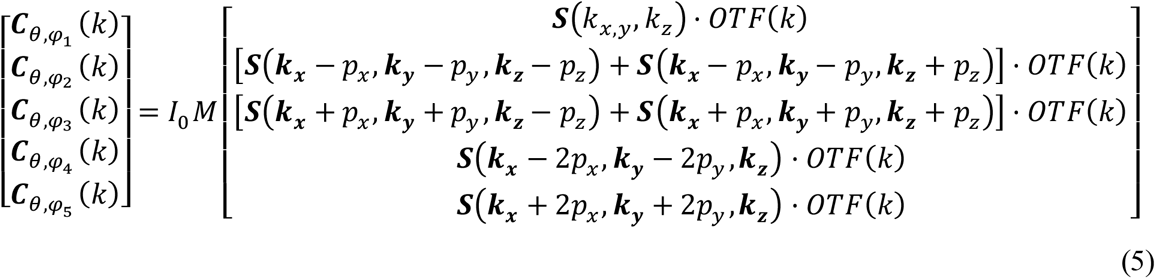

where 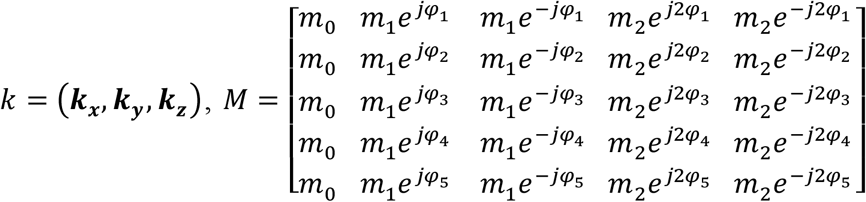 is the separation matrix, so the five bands ***B***_**0**_(*k****), B***_±1_(*k*), and ***B***_±2_(*k*) of 0^th^, ±1^st^, and ±2^nd^ can be expressed as:

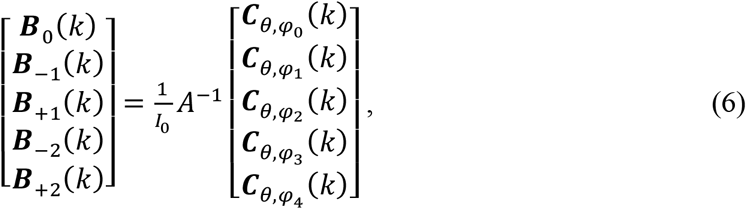

where [·]^−1^ is the symbol of the matrix inverse transform. Then the five components should be shifted into the correct spectrum position on the *xoy* plane, so the combined spectrum can be expressed as:

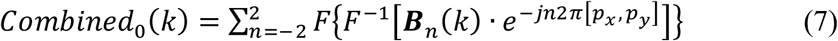

Where *F*[·] and *F*^−1^[·] denotes the 3D Fourier transform and Fourier inverse transform. Then, we use a notch filter^23^ and two steps of filters using Open-3DSIM to make the spectrum smooth and even and reduce the artifacts. The detailed steps are dismissed here and can be seen in Open-3DSIM^24^. When *Combined*_0_(*k*) is notched and filtered, it can be expressed as *Combined*_1_(*k*). Thus, the final 3DSIM result *S*_3*DSIM*_ (*x, y, z*) is:

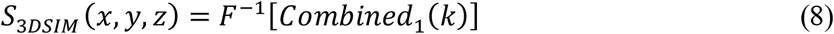

The detailed algorithm flow is shown in Fig. 2(c).

### 2.4 Open-source 3DSIM Hardware

Together with our recently published software of Open-3DSIM^24^, we have made all the hardware components and control mechanisms openly available in this study as a complementary open-source hardware counterpart. We have opened the detailed hardware devices in Supplementary S5 and the LabVIEW program in GitHub (https://github.com/Cao-ruijie/DMD-3DSIM-hardware) to control the sequence of DMD, sCMOS camera, EOM, and piezoelectric translation platform for reproduction and development of the 3DSIM system (Supplementary S6). Given the widespread utilization of DMD technology in digital light processing and its costeffectiveness as a consumer product, we anticipate that our initiatives will significantly support the community in devising diverse adaptations and iterations, capitalizing on the foundation established by our open hardware approach.

## 3 Results

To quantify the enhanced resolution and qualify the optical sectioning capability of our 3DSIM approach, we initially imaged 100 nm fluorescent beads used a raw-frame exposure time of 15 ms and the volume time of 6.75s (29 sections, 15 frames/section, 125 nm sectioning step). A comparison was made between the wide-field (WF) image and 3DSIM image as depicted in Fig. 3(a). In the magnified view of the region outlined by the blue rectangle, the 3DSIM image successfully resolves three adjacent beads that are indistinguishable in the WF image, highlighting a significant improvement in lateral resolution. Furthermore, the *xoz* cross-sectional images of the isolated beads at the bottom left of Fig. 3(a) demonstrate the enhanced optical sectioning capability of 3DSIM, effectively eliminating out-of-focus background both above and below the focal plane. The WF and 3DSIM intensity profiles of the fluorescent beads along the white line in the *xoy* and *xoz* planes are illustrated at the bottom right of Fig. 3(a). Additionally, image decorrelation method^25^ was performed to assess the resolution. The results revealed a lateral resolution of 263 nm in WF [Fig. 3(b)] and 133 nm in 3DSIM [Fig. 3(c)], showcasing a nearly 2-fold enhancement compared to WF. We also used PFSj^26^ on the images of the beads in Fig. 3(a) to evaluate the axial resolution of our system and found that 3DSIM can achieve the 2-fold resolution improvement in the axial direction from 601nm to 300m. Subsequently, dot-like samples of nuclear pore complex resembling fluorescent beads were imaged under the same acquisition timing settings. Fig. 3(c) showcases the WF and 3DSIM images of the nuclear pore complex within the cell nucleus. Similarly, 3DSIM improves the resolution in all three dimensions compared to WF and effectively eliminates defocus of the background, enabling a comprehensive interpretation of the entire nuclear structure.

**Fig. 3.**
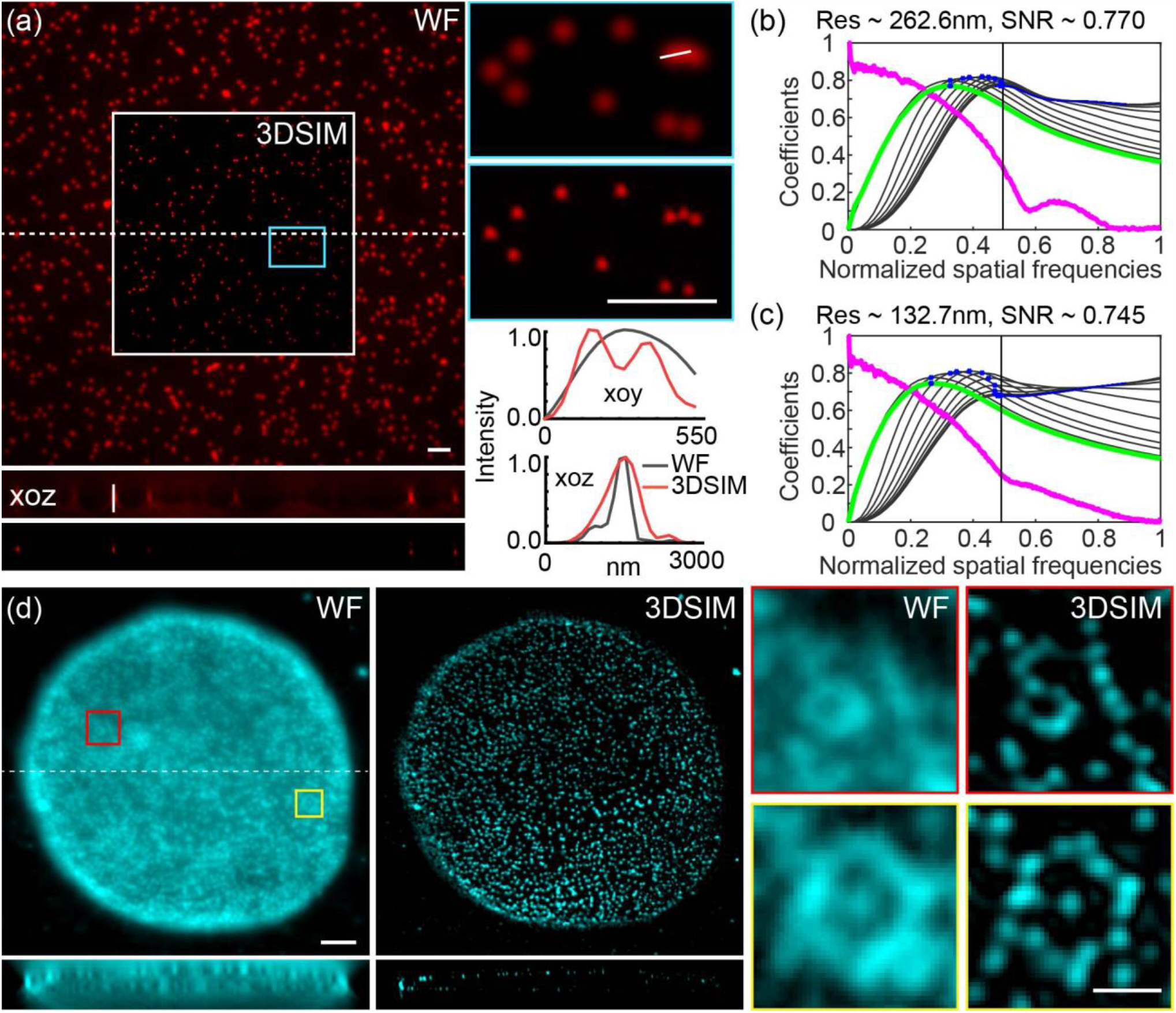
Resolution test of DMD-based 3DSIM system and super-resolution imaging of nuclear pore complex. (a) The WF and 3DSIM images of 100 nm fluorescent beads with the corresponding profiles of the intensity distribution (bottom right) along the magnified beads (top right) and the *xoz* view (bottom left) indicate the improved resolution in the *xoy* and *xoz* planes. (b-c) The quantitative analysis of decorrelation shows accurate WF and 3DSIM resolution in *xoy* planes. (d) The WF and 3DSIM images of nuclear pore complex of Cos-7 cells. (a ) 29 layers. (d) 29 layers. Scale bar: 2 μm in the whole field-of-view and 0.5 μm in the enlarged region-of-interest.

### 3DSIM and Polar-3DSIM imaging of subcellular organelles

We utilized 3DSIM to image more intricate subcellular structures with finely detailed three-dimensional features encompassed mitochondrial cristae and outer membranes, microtubule proteins, and actin filaments. Fig. 4(a) and Fig. 4(b) present WF and 3DSIM images showcasing the mitochondrial cristae and outer membrane, respectively. It can be seen 3DSIM image provides a distinct view of the intricate internal structure of the mitochondrial cristae while WF cannot in Fig. 4(a). Furthermore, the 3DSIM image of the mitochondrial outer membrane exhibits higher resolution compared to WF, with a reduced impact in the defocus background in Fig. 4(b). Furthermore, we conducted imaging of two linear subcellular structures, namely actin, and tubulin. Fig. 4(c) illustrates the imaging results for tubulin in COS7 where individual microtubules can be traced and observed throughout the entire cell volume, and many parallel microtubules can be distinctly resolved as separate entities. These findings demonstrate the superior capability of 3DSIM in revealing fine details and enhancing resolution for subcellular structures. In addition, we introduce polarization dimension^27-29^ on the *xoy* plane, namely polarized super-resolution 3DSIM imaging techniques (p3DSIM), to investigate the subcellular structure of actin filament in U2OS with unprecedented resolution and polarization sensitivity in Fig. 4(d). p3DSIM offers a clearer insight on the orientation and alignment of the actin filaments that the ensemble orientation of the dipoles is approximately parallel to the direction of the filaments, contributing to a deeper understanding of their functional organization within the cellular context. It’s noteworthy that with intensity calibration, the dipole orientation will be more accurately resolved by 3DSIM and Supplementary S7 shows the intensity calibration of different angles in our 3DSIM system. What’s more, we perfromed the comparison between WF, 2DSIM,and 3DSIM to demonstrate the advantage of 3DSIM in Supplementary Note 8, and the comparison between DMD-based 3DSIM (DMD-3DSIM) and SLM-based 3DSIM(SLM-3DSIM) in Supplementary Note 9.

**Fig. 4.**
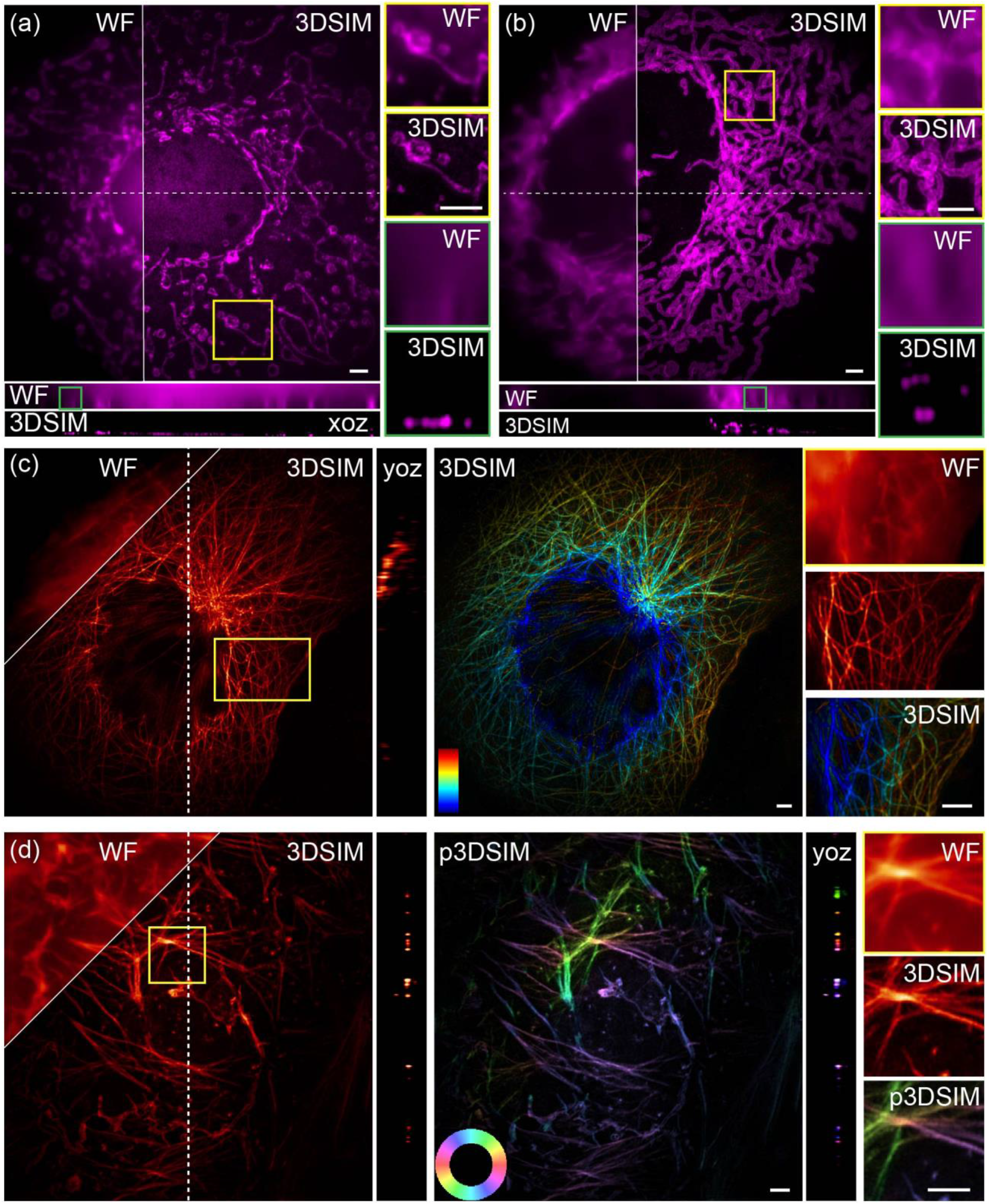
3DSIM reconstruction of subcellular structures. The maximum intensity projection images of (a) mitochondrial cristae, and (b) mitochondrial outer membrane with the corresponding magnified zone. (c) The maximum intensity projection images of tubulin in the form of a depth-color map. (d) Orientation imaging of actin filament and comparisons between WF, 3DSIM, and p3DSIM. The exposure times in (a-d) are 25 ms, 20 ms, 15 ms and 18 ms, respectively. (a) 29 layers. (b) 22layers. (c) 23 layers. (d) 25 layers. Scale bar: 2 μm.

**Fig. 5.**
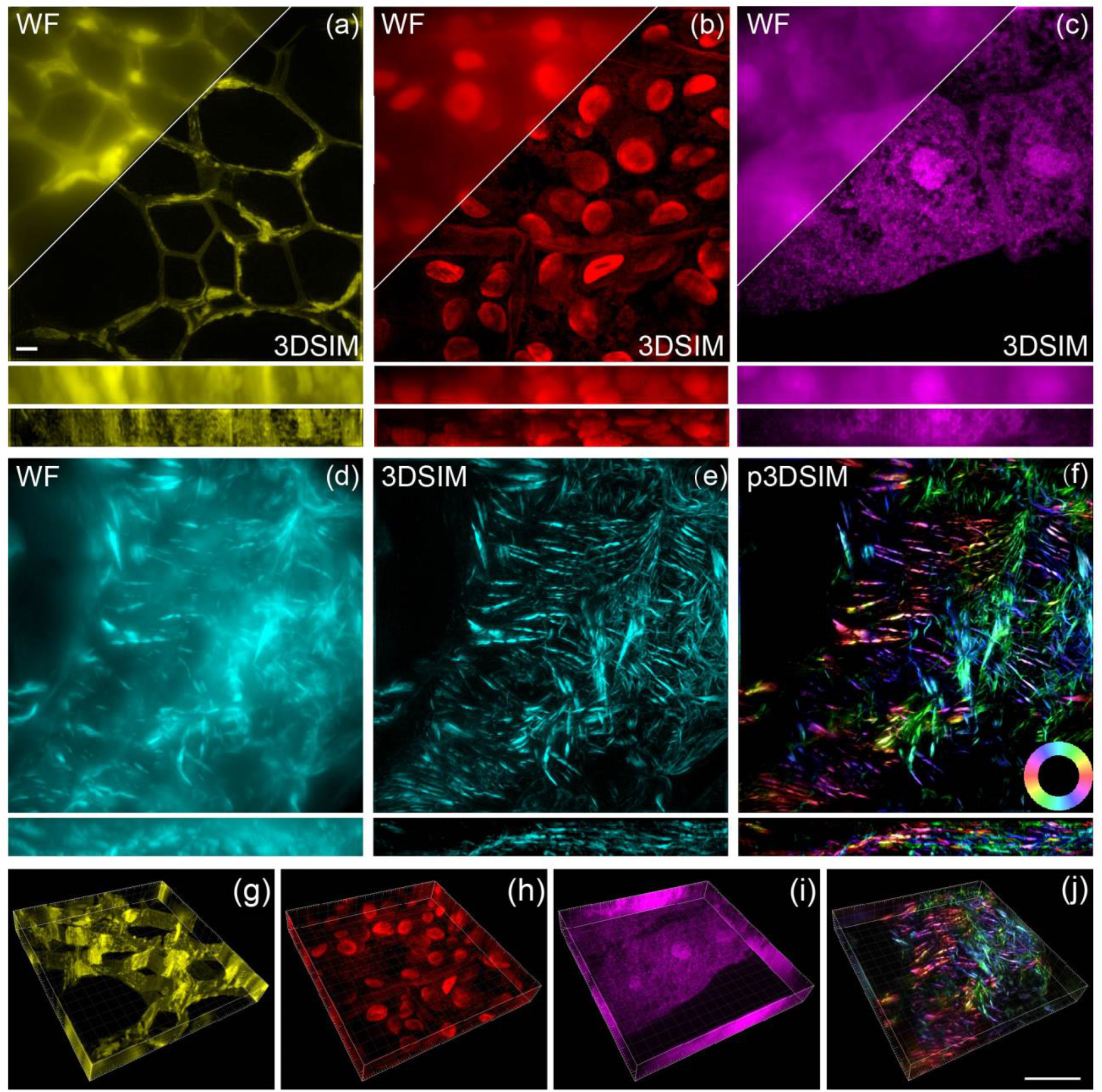
3DSIM reconstruction of plant and animal tissue samples. The maximum intensity projection (MIP) images of (a) cell walls in oleander leaves, (b) the hollow structures within black algal leaves, and (c) the periodic aggregation and dispersion of structures in the root tips of corn tassels. The corresponding 3D MIP images are respectively shown at the bottom. (d-f) The WF, 3DSIM, and p3DSIM MIP images of actin filaments in the mouse kidney tissue slice, with the corresponding *xoz* cross-sections a long the dashed line direction in (d). (g-j) The three-dimensional spatial distribution of (a-c) and (f). The exposure time in (a -d) are 20 ms, 15 ms, 25 ms and 18 ms, respectively. (a -d) 37 layers. Scale bar: 2 μm.

### 3D-SIM super-resolution of thick scattering specimen

We employed 3DSIM to image thick plant and animal tissues with a stronger defocus background. Fig. 6(a-c) presents the WF and 3DSIM results of oleander leaves, black algal leaves, and corn tassels, respectively. In the 3DSIM images, distinct features become apparent that were not observable in conventional WF images. These include the three-dimensional distribution of cell walls in oleander leaves, the hollow structures within black algal leaves, and the periodic aggregation and dispersion of structures in the root tips of corn tassels. In addition, we further analyzed the orientation of fluorescent dipoles within actin filaments in the mouse kidney tissue slice in Fig. 6(d). In the maximum projection image obtained from WF of the mouse kidney slices, the presence of out-of-focus scattered signals and the inability to distinguish in-focus information resulted in a less pronounced visualization of the filamentous structure. However, 3DSIM endows it with superior optical sectioning capabilities, presenting a clear representation of actin filaments, due to the higher resolution. Moreover, we utilized Polar-3DSIM to investigate again the alignment of fluorescent dipoles within actin filaments in 8.5 μm-depth mouse kidney tissue sections. Our findings demonstrated a prevailing parallel orientation of the dipoles relative to the filament axis. Additionally, Fig. 6(g)-(j) and Supplementary Note 10 show the spatial distribution of these samples in a three-dimensional manner, effectively highlighting the advantages of 3DSIM in enhancing resolution and removing the out-of-focus background. Furthermore, Supplementary Video 1 of black algal leaves and Supplementary Video 2 of actin filaments in the mouse kidney tissue slice accompany the manuscript, illustrating the three-dimensional reconstruction capabilities of 3DSIM, and providing a detailed visualization of angles of view.

**Fig. 6.**
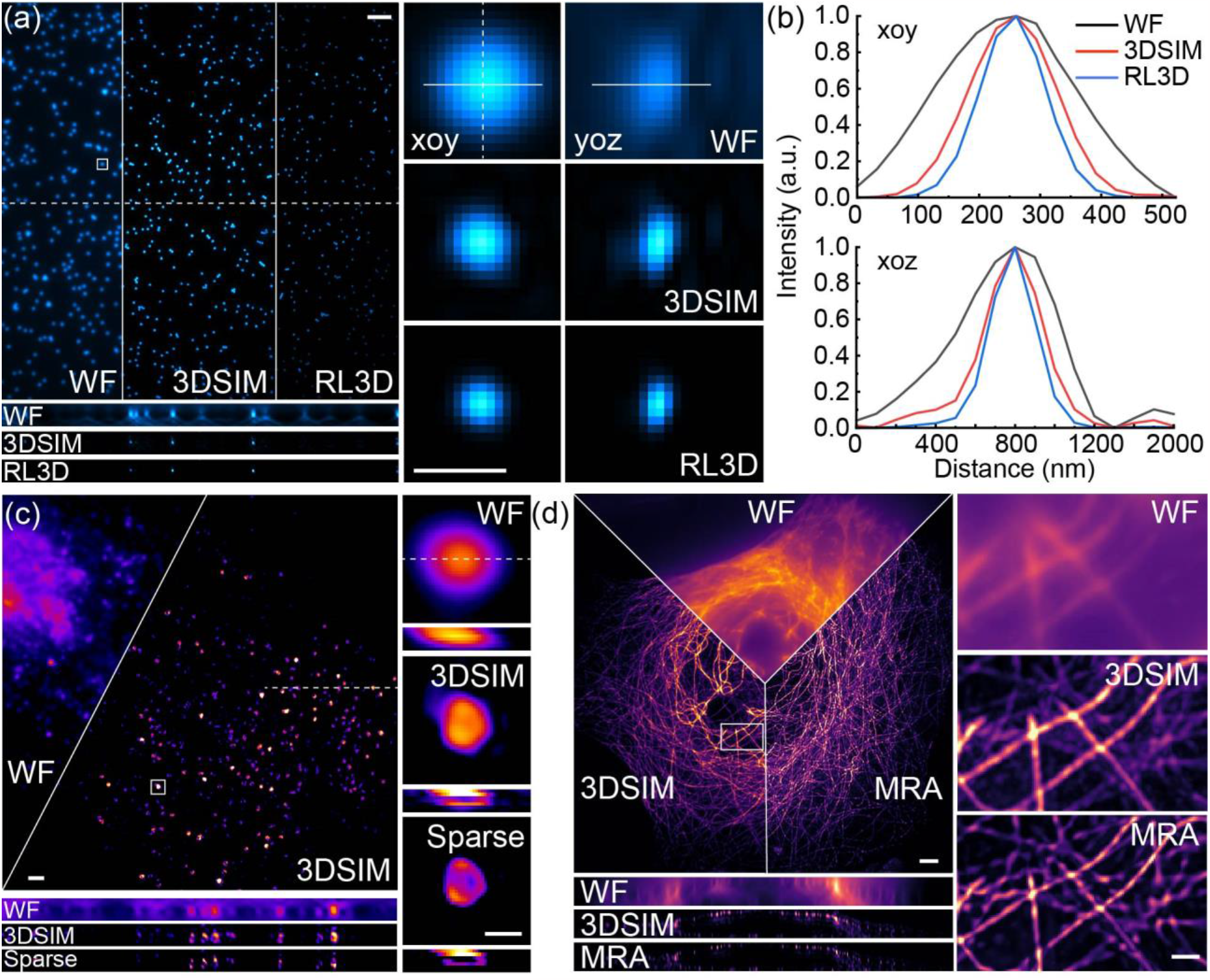
Deconvolution algorithms further improve the resolution of the 3DSIM system. (a) The WF, 3DSIM, and RL3D results of 100 nm beads in the 12^th^ layer with (b) the corresponding profile of the intensity distribution along the magnified bead, indicating the improved resolution in *xoy* and *xoz* planes. (c) The WF, 3DSIM, and Sparse deconvolution results of CCPs (labeled by recombinant anti-clathrin heavy chain antibody-Alexa Fluor 555) in COS7, where 3DSIM and Sparse deconvolution interpret the 3D hollow structure while WF cannot. (d) The WF, 3DSIM, and MRA deconvolution results of tubulin of Cos-7 cells, where MRA can further improve the 3D resolution of 3DSIM. The exposure times are 15 ms in (a), 25 ms in (c), and 15 ms in (d). (a) 24 layers. (c) 14 layers. (d) 22 layers. Scale bar: 2 μm in the whole field-of-view and 0.5 μm in the enlarged region-of-interest.

### Computational 3D deconvolution

Lastly, we use different 3D deconvolution algorithms to prove that computational super-resolution technology can further improve the 3D resolution of our 3DSIM approach. We use 3D Richardson Lucy (RL3D) deconvolution^30^ for the post-processing of the 3DSIM image of 100 nm beads as shown in Fig. 6(a), finding that RL3D can further improve 3D resolution compared with 3DSIM as the profiles in Fig. 6(b). We use Sparse deconvolution^31^ to do the post-processing of 3DSIM results of clathrin-coated pits (CCPs) in Fig. 6(c). Although 3DSIM can interpret the 3D hollow structure of CCPs, Sparse deconvolution can make the hollow structure clearer. This is in consistent with the physiological structure of CCPs^32^, and similar results were reported with 3D single-molecule localization microscopy^33^. Then we use MRA deconvolution^34^ for the processing of tubulin structure of low signal-to-noise ratio in Fig. 6(d). We can find that the artifacts of 3DSIM caused by the defocus and noise can be greatly suppressed using MRA, with an improved 3D resolution. These results show that our solution is suitable for many post-processing algorithms, which can further improve the resolution and quality of 3DSIM reconstruction.

## 4 Discussion and Conclusion

Firstly, due to the limitation of experimental condition (we do not have live cell incubation system in our custom-built microscope), our current DMD-based 3DSIM setup can only perform single-color imaging with fixed specimen. But we can further introduce other lasers to incident DMD at different angles to realize multi-color imaging without change on the post part of the system as referred to in our proposed four-color DMD-based SIM concept^35^. Meanwhile, regarding the potential applications of DMD-3DSIM for real-time and dynamic live cell imaging, it is better to equip the system with microscope body and live cell incubator, for spatial and temporal stability. And the fast switch of EOM and DMD exploits the potential to conduct the imaging of fast-moving living cells with the help of these improvement.

Secondly, in our DMD-based 3DSIM system, we conducted tests to measure the maximal data acquisition speed for single-layer 3DSIM with varying field-of-view (FOV) sizes in synchronous readout trigger mode (Supplementary Note 11). We can achieve the maximum 24 frame/s imaging when the FOV is 512×512 in 3DSIM modality, which depends largely on the readout and exposure time of the sCMOS because the switching time of DMD and EOM is several magnitudes higher than sCMOS. Furthermore, we performed 3DSIM imaging of tubulin in COS7 cells at the maximum illumination power with progressively decreasing exposure times (Fig. S6). However, when exposure times are further reduced (indicating faster imaging), the acquired raw data may exhibit inadequate signal-to-noise ratios, resulting in excessive reconstruction artifacts or even rendering the reconstruction impossible. Hence, we conclude several other factors which can influence the acquisition speed of multi-layer 3DSIM data in practical experiments even with the coordinated optimization of hardware and software. These include the characteristics of the illumination source (laser power and stability are crucial as they affect signal-to-noise ratios), sample characteristics (fluorescent samples with high and stable brightness perform better in fast imaging), and sample mounting and stability (any instability in the sample stage can lead to longer acquisition time).

Thirdly, although we have used various deconvolution algorithms to optimize the images (both Sparse and MRA deconvolution use the 2D OTF without considering the complex defocus background), advanced 3D algorithms^36, 37^ are still required to faithfully reconstruct and optimize low signal-to-noise ratio data acquired from high-speed live cell imaging.

In summary, we demonstrated our DMD-based 3DSIM system as a fast and reliable tool for subcellular structures, tissue architectures, and fluorescent dipole imaging with a 3D spatial resolution twice that of WF, and computational super-resolution algorithm can further improve the resolution of our system. Our system features the advantages of fast switching and accurate polarization-modulation, opening up a new avenue of exploration and providing invaluable tool for unraveling the complex mechanisms underlying fundamental biological processes.

## Acknowledgments

This work was supported by the National Key R&D Program of China (2022YFC3401100), National Natural Science Foundation of China (62335008, 62025501, 31971376, 92150301).

## Availability of data and materials

The data that support the findings of this study are available from the corresponding author on reasonable request. The LabVIEW program has been uploaded to https://github.com/Cao-ruijie/DMD-3DSIM-hardware.

