## Supplementary material for "High-speed Auto-Polarization Synchronization Modulation Three-dimensional Structured Illumination Microscopy"

### Supplementary materials for
### High-speed Auto-Polarization Synchronization Modulation Three-dimensional Structured Illumination Microscopy

**Yaning Li,[a,b,†] Ruijie Cao,[a,b,†] Wei Ren,[a,b] Yunzhe Fu,[a,b] Yiwei Hou,[a,b] Suyi Zhong,[a,b] Karl Zhanghao, [a,b] Meiqi Li,[a,b,*] Peng Xi[a,b,*]**

[a] Department of Biomedical Engineering, College of Future Technology, Peking University, No. 5 Yiheyuan Road, Beijing, China, 100871

[b] National Biomedical Imaging Center, Peking University, No. 5 Yiheyuan Road, Beijing, China, 100871

**S1. Comparison of three representative SIM methods with various polarization control schemes.**

In grating-based SIM, a linear polarizer is fixed on the rotatable table and rotates with the phase grating to make the polarization direction (s-polarization) of beams the same as structured illumination fringes, it can provide the best modulation depth of the illumination pattern. In FLC-SLM-based SIM and DMD-based SIM, zero-order vortex half-wave retarders[1] and pizza wave plates[2] are choices of the passive polarization control device of 2DSIM with the advantage of not requiring a timing sequence setting. In 3DSIM, the polarization state of the central zero-order beam also needs to be dynamically adjusted according to the direction of the loaded pattern images in FLC-SLM or DMD. This requirement stems from the necessity to maintain proper alignment between the polarization direction and the desired orientation of the interference fringes in the system. Therefore, in FLC-SLM-based 3DSIM, the polarization rotation active device can be implemented by combining two ferroelectric liquid crystal phase retarders with a quarter wave plate[3]. Similarly, in DMD-based 3DSIM, the polarization rotation device can be realized by combining one electro-optic modulator with a quarter wave plate. These configurations allow for precise control of the polarization state of all three beams. Table. S1 shows the comparison of three representative SIM methods with various polarization control schemes. We can see that DMD-based SIM with the electro-optic modulator has an ultrafast switching speed and the highest efficiency, which can generally suitable for both 2DSIM and 3DSIM imaging modes. So this is the reason for us to choose the DMD and EOM for our system.

**Table. S1** The comparison of three representative SIM methods with various polarization control schemes.

| SIM setup Category | Polarization control device | Classification | Switching | Efficiency (%) | Configuration |
|---|---|---|---|---|---|
| Grating-based SIM | polarizer | passive | -- | ~90 | 2DSIM, 3DSIM |
| FLC-SLM-based SIM | Zero-order vortex half-wave retarder | passive | -- | ~90 | 2DSIM |
| | Pizza wave plate of six-zone | passive | -- | ~40 | 2DSIM |
| | Ferroelectric liquid crystal phase retarders | active | fast | ~75 | 2DSIM, 3DSIM |
| DMD-based SIM | Zero-order vortex half-wave retarder | passive | -- | ~90 | 2DSIM |
| | Pizza wave plate of six-zone | passive | -- | ~40 | 3DSIM |
| | Electro-optic modulator | active | ultrafast | ~90 | 2DSIM, 3DSIM |

**S2. The polarization extinction ratio of DMD-based SIM with three polarization control schemes.**

In DMD-based SIM, the utilization of a zero-order vortex half-wave retarder or pizza wave plate of six-zone is limited to 2DSIM applications with a relatively low extinction ratio at the back focal plane of the objective. To overcome these limitations, an alternative approach employing an EOM for polarization modulation has been proposed. This method offers several advantages as it applies to both 2DSIM and 3DSIM and provides a higher extinction ratio at the back focal plane of the objective. By incorporating the EOM-based polarization modulation scheme into the DMD-based SIM system, we can achieve high-quality imaging of subcellular structures with increased contrast and fidelity. Comparisons of DMD-based SIM with zero-order vortex half-wave retarder, pizza wave plate of six-zone, and EOM are shown in Table. S2-4 we can see that EOM achieved the best extinction ratio at the back focal plane of the objective, providing the best contrast of interferometric fringes for better SIM reconstruction.

**Table. S2** Polarization extinction ratio of DMD-based SIM with Zero-order vortex half-wave retarder.

| Beam | Zero-order vortex half-wave retarder | Focal plane of objective |
|---|---|---|
| Direction1, +1 | 20.00 | 83.50 |
| Direction1, -1 | 190.02 | 20.83 |
| Direction2, +1 | 17.89 | 115.88 |
| Direction2, -1 | 89.60 | 18.89 |
| Direction3, +1 | 72.21 | 20.35 |
| Direction3, -1 | 9.19 | 7.34 |

**Table. S3** Polarization extinction ratio of DMD-based SIM with Pizza wave plate of six-zone.

| Beam | Pizza wave plate of six-zone | Focal plane of objective |
|---|---|---|
| Direction1, +1 | 31.05 | 25.59 |
| Direction1, -1 | 26.31 | 23.66 |
| Direction2, +1 | 84.00 | 62.5 |
| Direction2, -1 | 112.31 | 29.7 |
| Direction3, +1 | 75.03 | 23.88 |
| Direction3, -1 | 38.55 | 72.14 |

**Table. S4** Polarization extinction ratio of DMD-based SIM with EOM.

| Beam | Style name | Focal plane of objective |
|---|---|---|
| Direction1, +1 | 678 | 226.27 |
| Direction1, -1 | 689 | 231.59 |
| Direction2, +1 | 336.5 | 92.86 |
| Direction2, -1 | 331 | 86.70 |
| Direction3, +1 | 650 | 203.42 |
| Direction3, -1 | 659 | 212.37 |

**S3. Construction of 3DSIM System.**

To achieve high-quality reconstructions in 3DSIM, it is crucial to generate an optimal illumination pattern with high modulation contrast in the optical setup. The generation of the pattern involves the interference of three coherent beams, which needs equal intensity and polarization for three beams. In previous experimental experience[4], the grating was specially designed so that the intensity of order 0 was 70–80% of that of the central order, this unequal intensity ratio was picked to slightly strengthen the high-frequency components. Therefore, we set DMD under the blazed grating condition[1], use a custom-built spatial mask in Fig. 1(a), and utilize an EOM with the best polarization extinction ratio to maintain these conditions ensuring complete destructive interference, resulting in maximum modulation contrast. By carefully controlling the intensity and polarization of the interfering beams, we can maximize the modulation contrast and enhance the quality of the 3DSIM reconstructions.

#### S4. The synchronization sequence of high-speed 3DSIM System.

During the data acquisition process in 3DSIM, several operations meeting the time sequence of the synchronization are required. This involves switching patterns (five pattern phases spaced evenly at intervals of 2π/5, three pattern orientations separated by 60 degrees) loaded on the DMD to generate the structured illumination stripes, adjusting the polarization state of the illuminating beam by varying the voltage applied to the electro-optic modulator, precisely controlling the axial movement using the piezo translation stage, and accurately triggering of camera exposure and readout. In our system, DMD has a minimum exposure time of 105 μs, thanks to the high switching speed of the DMD's digital controller (DLPC900), capable of switching 1-bit binary patterns at a rate of up to 9523 Hz. Additionally, the EOM exhibits a high polarization modulation speed of 250 kHz. Therefore, excluding the response time of the piezo translation stage for axial movement, the imaging speed of our constructed 3DSIM system is solely limited by the performance of current-generation sCMOS cameras. The synchronous readout trigger mode of the sCMOS is used for continuous imaging. In this mode, the camera ends each exposure, starts the readout, and starts the next exposure at the edge of the input trigger signal (rising edge) at the same time. Fig. S1 shows the time synchronization sequence of our 3DSIM approach, with which our 3DSIM can run at 112 frames per second with a 90 × 90 pixel (5.85 × 5.85 micron) field of view. It is worth noting that the imaging speed of 3DSIM described here holds great potential for real-time and dynamic imaging applications. The rapid acquisition of volumetric data sets opens up possibilities for investigating dynamic processes and studying time-dependent cellular events with high temporal resolution. Moreover, the compatibility of our system with the latest-generation sCMOS cameras ensures the capture of high-quality images with improved sensitivity and reduced noise levels.

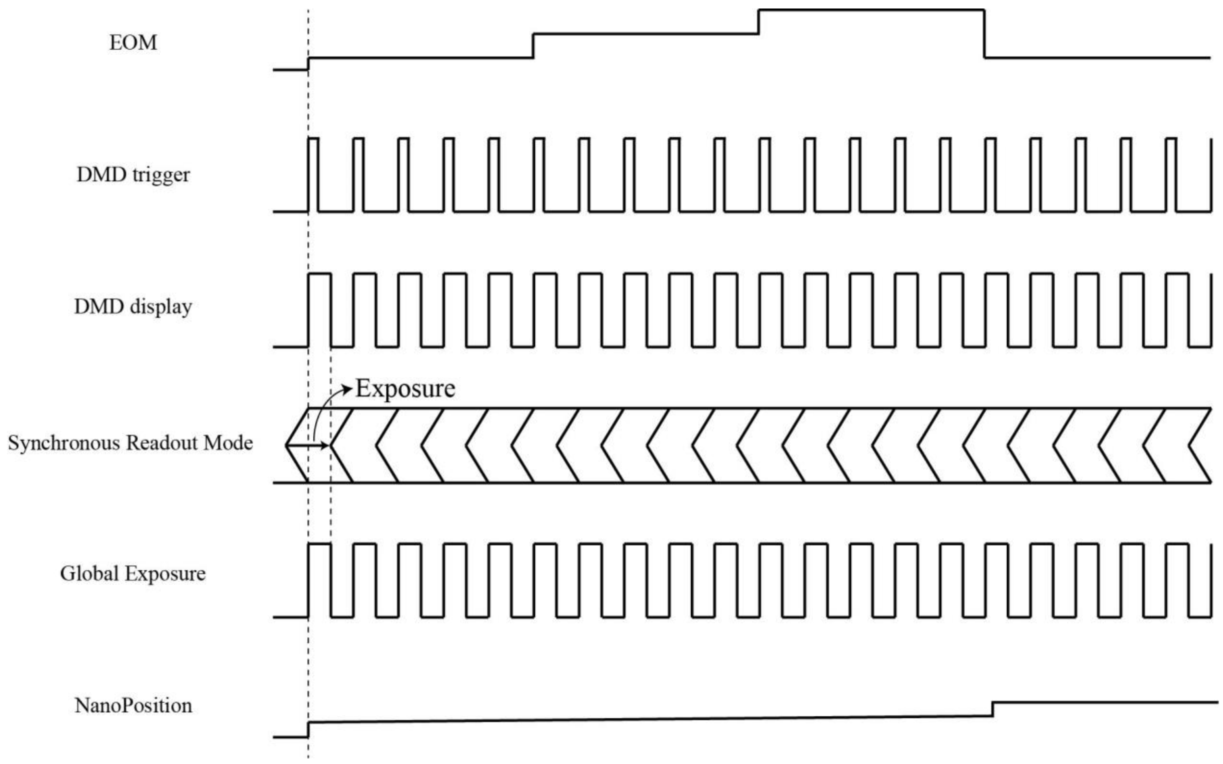

**Fig. S1** Time sequence of the synchronization of DMD-based 3DSIM. Trigger signal of the EOM (row1) and DMD (row2). The DMD is preloaded with a series of patterns (row 3). The sCMOS camera operates in synchronous readout mode, with an 87.7 µs delay after receiving the rising edge (row 4). The output from the DMD triggers the sCMOS camera. Trigger signal of the Nano-translation platform (row 6).

#### S5. Open-source hardware devices of high-speed auto-polarization 3DSIM system.

**Table. S5** Open-source hardware devices list of high-speed auto-polarization 3DSIM system.

| Optical element | Vendor | Part number | Quantity | Unit Price (CNY) |
|---|---|---|---|---|
| Laser | Changchun New Industries Optoelectronics Technology | MGL-FN-561-200mW | 1 | 28000.00 |
| Glan-Taylor polarizing prism | LBTek | GTP10-A | 1 | 5629.00 |
| Electro-optic modulator | Conoptics | M370 | 1 | 99850.00 |
| Voltage differential amplifier | Conoptics | 302RM | 1 | 54650.25 |
| Quarter wave plate | Union Optic | wpa4420-450-650 | 1 | 2850.00 |
| Beam expander | Thorlabs | GBE10-A | 1 | 5,647.24 |
| Digital micro-mirror device | Texas Instruments | DLP6500 0.65 1080p MVSP Type A DMD | 1 | 15394.18 |
| Lens | Thorlabs | AC254-300-A-ML | 3 | 995.45 |
| Spatial mask | custom-built | ---- | 1 | 200.00 |
| Objective lens | Nikon | CFI Apochromat TIRF 100XC 1.49NA Oil | 1 | 108452.00 |
| Dichroic mirror | Chroma | ZT405-415/488/561/640-phaseR-UF | 1 | 4950.00 |
| Tube lens | Thorlabs | TTL200-A | 1 | 4,578.70 |
| Emission filter | Chroma | ET620/60m | 1 | 2,750.00 |
| Camera | Hamamatsu | ORCA-Flash4.0 V2 | 1 | 130748.56 |
| Water chiller | Coolium Instruments | LX-150 | 1 | 4500.00 |
| Silver mirrors | Thorlabs | PF10-03-P01 | 13 | 490.66 |
| Data AcQuisition Device | National Instruments | PCIe-6738 | 1 | 19105.36 |
| Sum | ---- | ---- | ---- | 496670.22 |

Note：

1、 For users who want to achieve better resolution and visualize fixed samples, shorter-wavelength laser sources of 360 nm (UV-FN-360 nm, Changchun New Industries Optoelectronics Technology, 175,000.00 CNY), the DMD (V-7001 UV, ViALUX, 122,000.00 CNY) and the objective lens (CFI SR HP Plan Apo Lambda S 100XC Sil, Nikon, 137,446.48 CNY) suitable for ultraviolet illumination are recommended.

2、 For users who want to obtain greater imaging depth, longer-wavelength laser sources of 976 nm (TH97-1050-7, Taizhou Tonghe Laser Technology, 11,600 CNY) and the DMD (AUT-V-650L NIR, ViALUX, 86,000.00 CNY) the objective lens (CFI SR HP Plan Apo Lambda S 100XC Sil, Nikon, 137,446.48 CNY) suitable for near-infrared illumination are preferred.

**S6. Open-source LabVIEW program of high-speed auto-polarization 3DSIM system.**

We independently developed the 3DSIM synchronization control program based on LabVIEW and its open-source program is linked to GitHub (https://github.com/Cao-ruijie/DMD-3DSIM-hardware). The main program covers five synchronous analog outputs, among which the analog output AO0 can be used for individual testing of each active device, the analog output AO2 is used for external trigger control of DMD, the analog output AO4 is used for external trigger control of piezoelectric translation platform, and the analog output AO6 is used for external trigger control of sCMOS camera. The analog output AO8 is used for external trigger control of the electro-optic modulator. The analog output of different channels is synchronized and can be monitored in the real-time output waveform. Fig.S2 shows the layout of the control panel of our LabVIEW program.

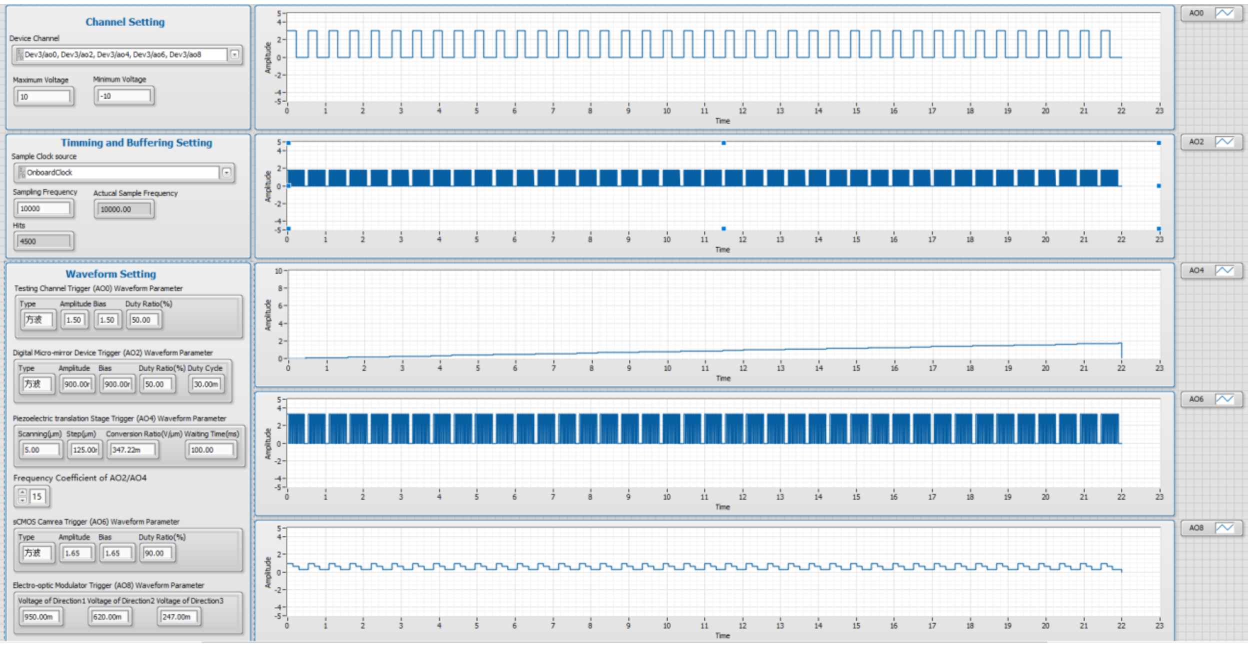

**Fig. S2** The layout of the control panel of the LabVIEW program of the high-speed auto-polarization aynchronization 3DSIM system.

Users need to complete three steps to use the open-source LabVIEW program. First, the user needs to select the correct analog output channel in the “Channel Setting” section according to the actual hardware connection interface. Secondly, the user needs to set the trigger voltage and trigger period of DMD, the “conversion ratio” of voltage/ motion range (V/μm) and waiting time of the piezoelectric translation platform, the trigger voltage of the sCMOS camera, and the three trigger voltages corresponding to the three directions of linearly polarized light generated by the electro-optic modulator in the “waveform setting” part according to the external trigger requirements of the specific hardware. Finally, the synchronization control of the 3DSIM system can be realized by running the software.

#### S7. Intensity calibration of dipole orientation imaging.

We have conducted the intensity calibration to make the interpretation of diploe orientation more accurate. We use the dense beads (Thermo Fisher Scientific, diameters of 100 nm) prepared in a fixed slide. The beads were imaged by our system and excited by light from 3 angles and 5 phases at each angle. We sum up the 5 images of phases in each angle to form the WF image with different angles as shown in Fig. S3(a) as Angle1, Angle2, and Angle3. Then we use the poly55 method to interpret the intensity map. Then we can get calib1=Angle2/Angle1, calib2=Angle3/Angle1 as shown in Fig. S3(b). calib1 and calib2 are used to correct the WF images of three angles to correct the polarization map.

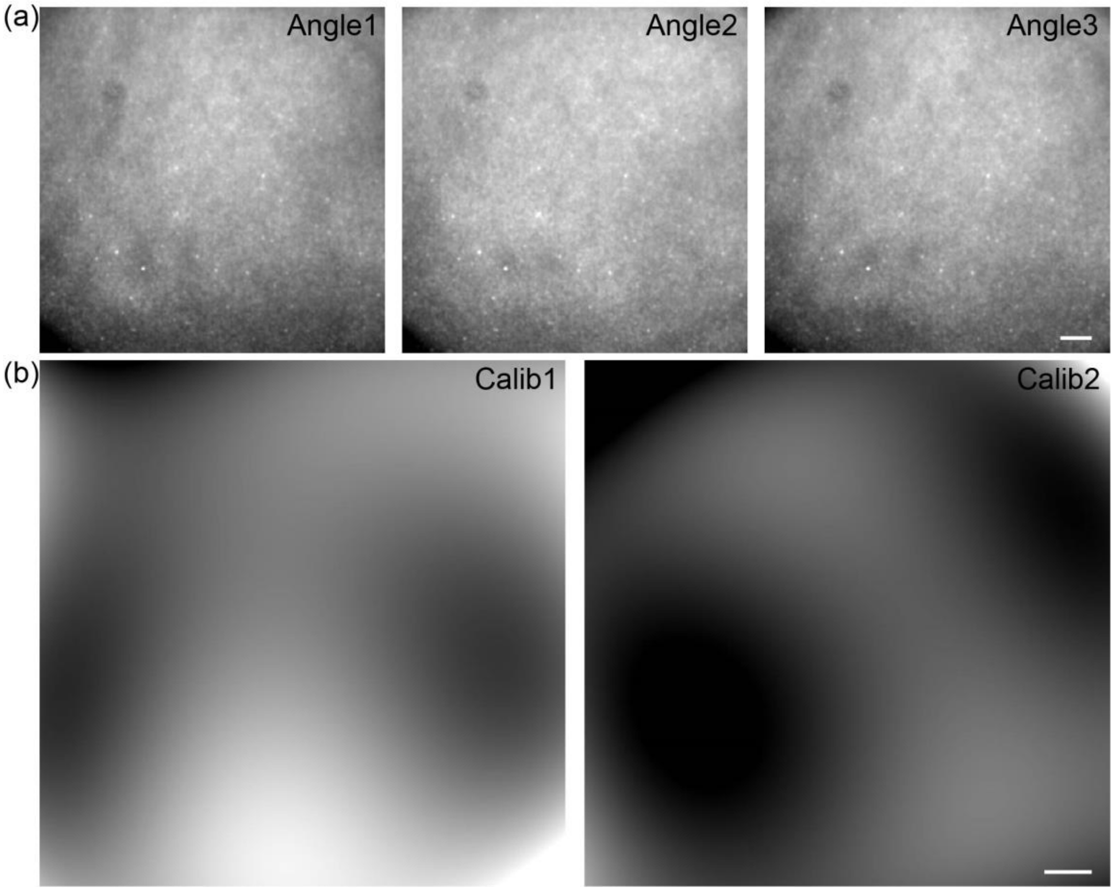

**Fig. S3** The intensity calibration of different angles in our system. (a) Images of beads in three excitation angles of Angle1, Angle2, and Angle3, (b) The calibration map of Calib1 and Calib2. Scale bar: 6 μm.

#### S8. The actin filament in U2OS cell imaged with WF, 2DSIM, and 3DSIM modality.

Our DMD-based 3DSIM microscope is currently capable of easily achieving WF, 2DSIM, and 3DSIM imaging simultaneously and we have added detailed comparison in Fig. S4 presents imaging results of actin filaments in U2OS cells under WF, 2DSIM, and 3DSIM modalities. It can be seen that 2DSIM has point-shaped artifacts due to the incomplete elimination of defocused background and severe artifacts in axial plane due to the low modulation on defocus plane. However, 3DIM can reduce the artifacts in both xoy and xoz plane, and achieved double resolution in axial plane compared with 2DSIM. And compared to the WF modality, 3DSIM significantly enhances both lateral and axial resolution.

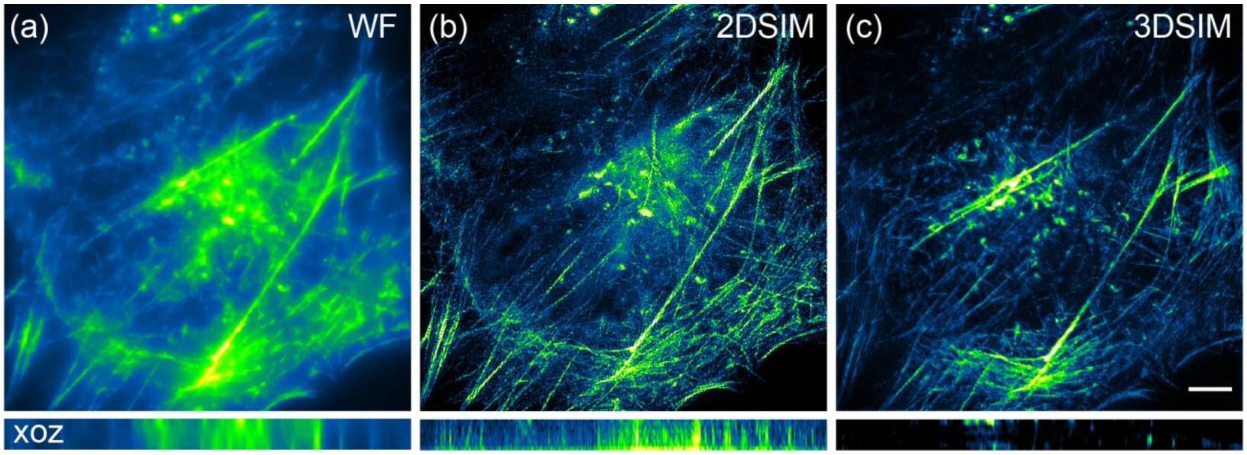

**Fig. S4** The actin filament in U2OS cell imaged with WF, 2DSIM, and 3DSIM modality in the same FOV. (a) WF. (b) 2DSIM. (c) 3DSIM. Scale bar: 4um. 10 layers.

#### S9. The comparison between DMD-3DSIM and SLM-3DSIM.

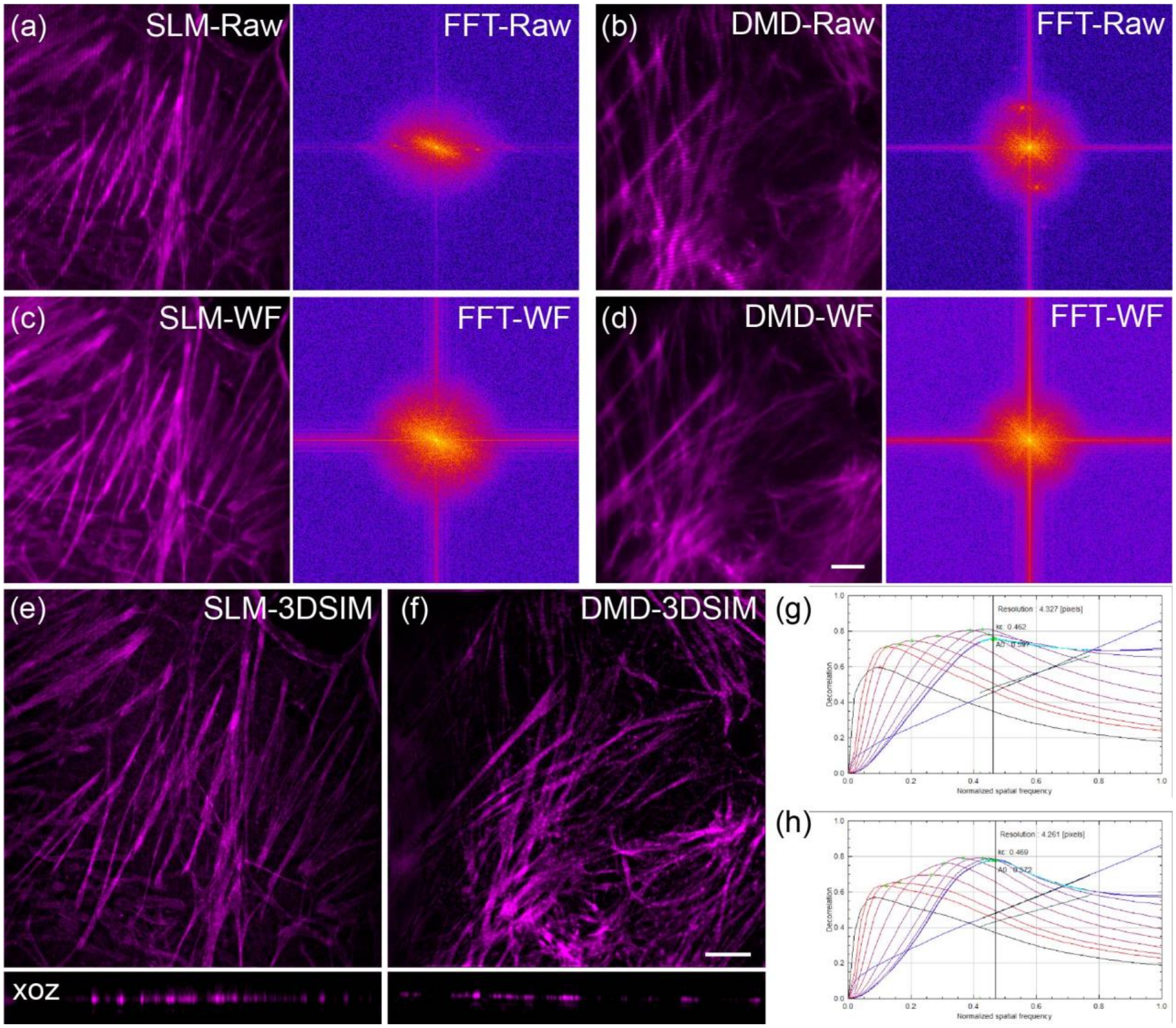

**Fig. S5** The comparison of optical modulation patterns and imaging results between DMD and SLM devices in 3DSIM. (a) One raw image of actin filament in U2OS and its fourier spectrum of SLM-based 3DSIM. (b) One raw image of actin filament in U2OS and its fourier spectrum of DMD-based 3DSIM. (c) The WF image calculated from a layer of 15 raw images of SLM-based 3DSIM data with the same FOV of (a). (d) The WF image calculated from a layer of 15 raw images of DMD-based 3DSIM data with the same FOV of (b). (e) The 3DSIM super-resolution image of SLM-based 3DSIM system with the same FOV of (a) and (c). (f) The 3DSIM super-resolution image of DMD-based 3DSIM system with the same FOV of (b) and (d). (g) The quantitative analysis of decorrelation shows accurate 3DSIM resolution of (e). (h) The quantitative analysis of decorrelation shows accurate 3DSIM resolution of (f). 21 layers in (e). 29 layers in (f). Scale bar: 4um.

We performed 3DSIM imaging using our laboratory-built SLM-based 3DSIM microscope and DMD-based 3DSIM microscope, respectively (Fig. S5). By loading precisely generated binary stripe patterns on the SLM and DMD, we obtained 3DSIM raw images with clear pattern, both of which exhibited distinct ±1 and ±2 order Fourie domain (Fig. S5(a), (b)). Through the averaging of 15 single-layer 3DSIM raw images, we acquired uniform wide-field images without pattern in spatial domain and without modulation point in Fourier domain (Fig. S5(c), (d)), demonstrating that the reproducibility and precise control of the light field, whether using DMD or SLM, can meet the requirements of 3DSIM imaging. Satisfactory super-resolution results were achieved for both the SLM-based 3DSIM microscope and the DMD-based 3DSIM microscope through the reconstruction using the open 3DSIM algorithm (Fig. S5(e), (f)).

In practical experiments, when choosing an objective with the same numerical aperture and laser excitation wavelength, the maximum achievable resolution is determined by the period of the interference pattern generated by the modulation structured illumination, and this interference pattern's period depends on the angle between the positive and negative first-order diffraction beams produced by the SLM or DMD. This angle is primarily determined by the period of the binary stripes loaded onto the SLM or DMD. When the stripe period loaded onto the SLM or DMD becomes smaller, the angle between the positive and negative first-order diffraction beams produced by diffraction increases. The angle can be very high with the participation of lens and small stipe period, but it is limited by diffraction limit. This means the maximum resolution enhancement is limited to twice because of diffraction limit rather than the binary strips loaded by DMD or SLM. So, the maximum resolution has no difference between DMD and SLM when wavelength and objective is fixed. In the experiments depicted in Fig. S5(e) and (f), we employed the same 561 nm wavelength laser as the excitation source and the same objective lens with the numerical aperture of 1.49. By loading binary stripe patterns with different periods onto both the SLM and DMD, we achieved a comparable enhancement in resolution of 140.6 nm and 138.5 nm respectively in SLM-SIM and DMD-SIM for the identical actin sample structure, as shown in Fig. Fig. S5(g) and (h).

**S10. The wide-field and 3DSIM volume comparison of oleander leaves, black algal leaves, and root tips of corn tassels.**

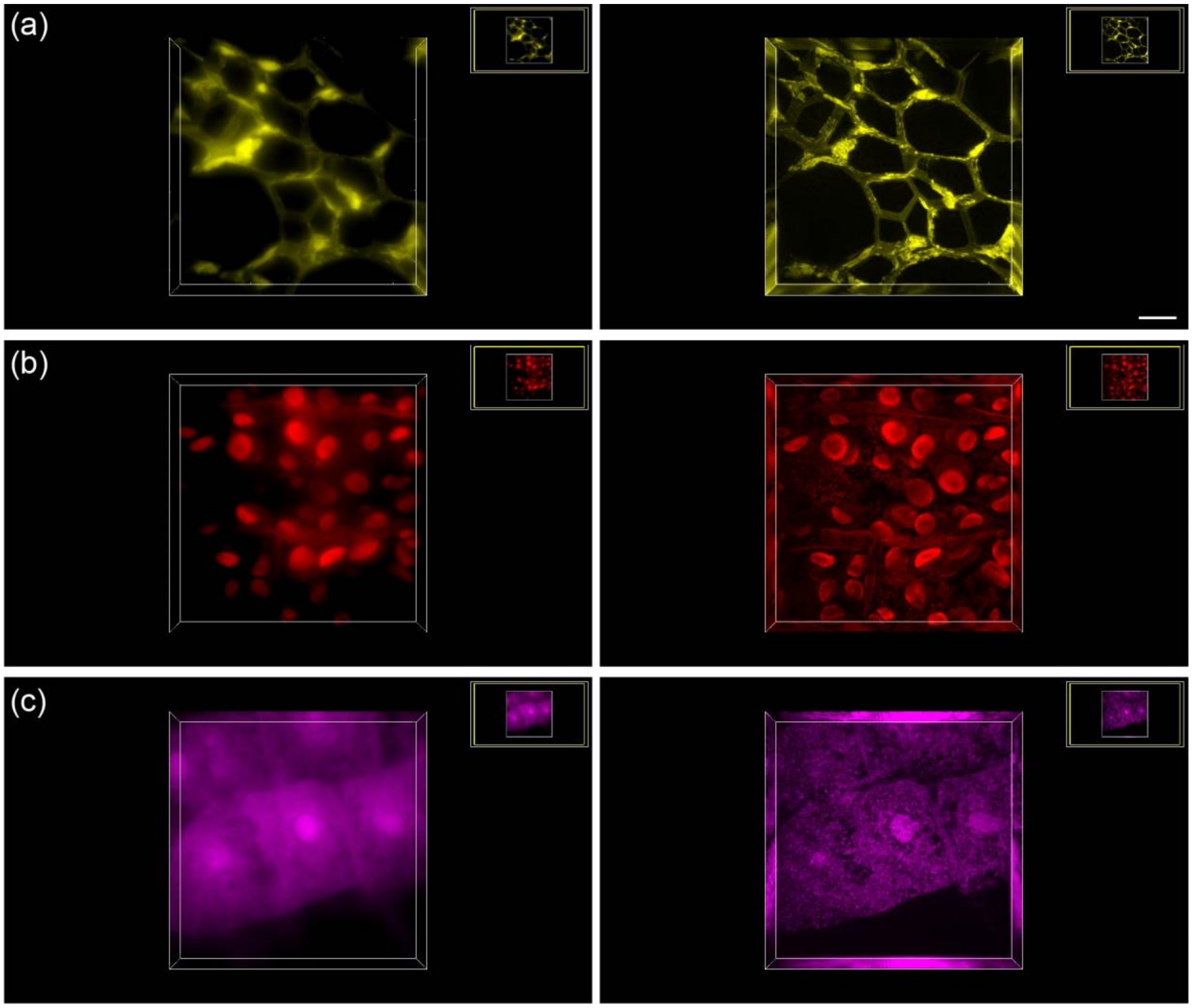

**Fig. S6** The three-dimensional display of (a) the cell walls in oleander leaves, (b) the hollow structures within black algal leaves, and (c) the periodic aggregation and dispersion of structures in the root tips of corn tassels with wide-field and 3DSIM images. Scale bar: 5 μm.

#### S11. The discussion about the imaging speed of DMD-based 3DSIM.

**Table. S6** The frame rate in synchronous readout trigger mode with USB 3.0 at standard scan.

| Field-of-View | Acquisition Fame Rate (frame/s) | Reconstruction Frame Rate (frame/s) |
|---|---|---|
| 8 × 512 | 4664 | 310 |
| 64 ×512 | 2052 | 136 |
| 128 × 512 | 1251 | 83 |
| 256 ×512 | 702 | 46 |
| 512 ×512 | 374 | 24 |
| 1024 ×512 | 193 | 13 |

We performed 3DSIM imaging of tubulin in COS7 cells at the same illumination power with the raw-frame exposure times of 45 ms, 15 ms, and 5 ms within the same FOV (The volume imaging duration was 8.2 s, 2.8 s, and 1.0 s, respectively) as shown in Fig. S6.

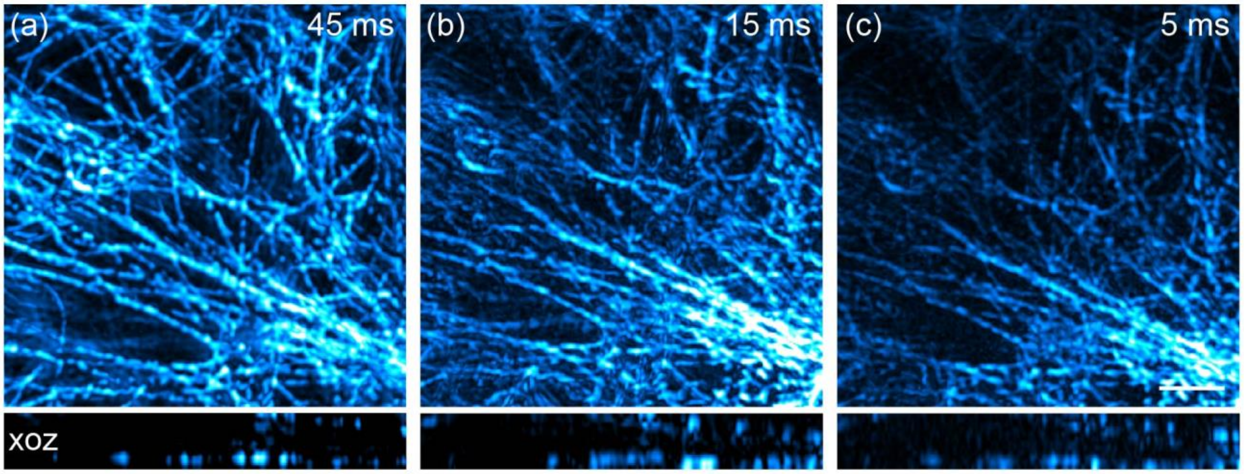

**Fig. S7** Tubulin in COS7 cell. The exposure times of (a-c) are 45 ms, 15 ms, and 5 ms, respectively. Scale bar: 4um. 12 layers.

#### S12. Video 1 | The three-dimensional rotation display of black algal leaves with 3DSIM.

#### S13. Video 2 | The three-dimensional rotation display of actin filaments in the mouse kidney tissue slice with p3DSIM.
